# Gut bacterial metabolite Urolithin A (UA) mitigates Ca^2+^ entry in T cells by regulating miR-10a-5p

**DOI:** 10.1101/255588

**Authors:** Shaqiu Zhang, Tamer Al-Maghout, Hang Cao, Lisann Pelzl, Madhuri S Salker, Anchun Cheng, Florian Lang, Yogesh Singh

## Abstract

The gut microbiota influences several biological functions including immune response. Inflammatory bowel disease is favourably influenced by consumption of several dietary natural plant products such as pomegranate, walnuts and berries containing polyphenolic compounds such as ellagitannins and ellagic acid. The gut microbiota metabolises ellagic acid leading to formation of bioactive urolithins A, B, C and D. Urolithin A (UA) is the most active and effective gut metabolite and acts as a potent anti-inflammatory and anti-oxidant agent. However, how gut metabolite UA affects the function of immune cells remained incompletely understood. T cell proliferation is stimulated by store operated Ca^2+^ entry (SOCE) resulting from stimulation of Orai1 by STIM1/STIM2. We show here that treatment of murine CD4^+^ T cells with UA (10 µM, 3 days) significantly blunted SOCE in CD4^+^ T cells, an effect paralleled by significant downregulation of Orai1 and STIM1/2 transcript levels and protein abundance. UA treatment further increased miR-10a-5p abundance in CD4^+^ T cells in a dose dependent fashion. Overexpression of miR-10a-5p significantly decreased STIM1/2 and Orai1 mRNA and protein levels as well as SOCE in CD4^+^ T cells. UA further decreased CD4^+^ T cell proliferation. Thus, bacterial metabolite UA up-regulates miR-10a-5p thus interfering with Orai1/STIM1/STIM2 expression, store operated Ca^2+^ entry and proliferation of murine CD4^+^ T cells.

## INTRODUCTION

Polyphenolic compounds modify inflammation, angiogenesis, drug and radiation resistance (1). Ellagitannin-rich food products and medicinal plants favourably influence inflammatory disease (2). *In vivo* studies from animal models indicate that ellagitannin-containing food products can be especially effective in modulating intestinal inflammation (3). The administration of pomegranate, raspberry, strawberry and almond preparations was shown to ameliorate the histological symptoms of chemically induced inflammation in gut mucosa, which was usually accompanied by decreased infiltration of immune cells, reduced overexpression of pro-inflammatory factors and the inhibition of inflammation associated molecular pathways (3–6). The bioavailability of ellagitannins and ellagic acid is, however, rather limited and the substances are metabolized by the gut microbiota yielding bioactive molecules including urolithins that are more readily absorbed than the original polyphenols (7). Urolithins circulate in plasma as glucuronide and sulphate conjugates at concentrations in the range of 0.2 – 20 µM (7). The characterization of the metabolites produced from polyphenols by gut microbiota is of clinical interest due to their antioxidant, anti-inflammatory and antiestrogenic activity (4). Thus, gut metabolites could modify immune cells including the adaptive immune T cells (8–11).

T-cells are activated by increase of intracellular Ca^2+^ (12, 13). T-cell receptor (TCR) engagement leads to activation of different signal transduction pathways that cause a rapid release of Ca^2+^ from the endoplasmic reticulum (ER) (14–17). In the resting state of T cells, Ca^2+^ is stored in the ER where it is sensed by stromal cell-interaction molecule (STIM) 1 and 2 proteins (18). Activation of the T cell receptor results in the production of inositol triphosphate (IP_3_), which binds to IP_3_ receptors on the ER and triggers the release of Ca^2+^ into the cytosol (12). The emptying of the intracellular Ca^2+^ stores is followed by store operated Ca^2+^ entry (SOCE) (12, 19–22) accomplished by activation of calcium release-activated calcium (CRAC) channel protein Orai1 by the ER Ca^2+^ sensors STIM1/2. Ca^2+^ influx through Orai1 in T cells depends on a negative membrane potential that provides the electrical driving force for Ca^2+^ entry (12, 21, 23–25). Ca^2+^ entry is required for full triggering of T-cell activation and proliferation, which involves expression of a large number of activation-associated genes (25).

T cell activation is modified by several microRNAs (miRNAs), which are post-transcriptional gene regulators (26–29). A recent study suggested that absence of miRNAs processing enzyme *dicer* inhibits the Ca^2+^ influx in naïve or activated T cells (30). Polyphenols such as green tea induces miR-15b which negatively affects the influx of Ca^2+^ inside the cells by regulating the Ca^2+^ sensing proteins STIM2 and Orai1 (31). However, whether gut metabolites such as UA influence miRNAs thus regulating the physiological functions of CD4^+^ T cells remained elusive.

In this study, we found a completely novel role of UA in the regulation of miRNAs expression, SOCE and proliferation of murine CD4^+^ T cells. Our results suggest that in CD4^+^ T cells, UA downregulates the expression of Orai1 and STIM1/2 thus compromising SOCE. It is further shown that UA upregulates expression of miR-10-5p in a dose-dependent manner, which in turn downregulates Orai1 and STIM1/2 transcript and protein levels thus blunting SOCE. Moreover, UA treatment decreased proliferation of CD4^+^ T cells. Thus, the present observations uncover a novel action of UA, i.e. the upregulation of miR-10a-5p with subsequent downregulation of store operated Ca^2+^ influx in CD4^+^ T cells.

## RESULTS

### UA attenuates store operated Ca^2+^ entry (SOCE)

Orai1 channels are recruited after being stimulated by STIM1/2, accomplishing SOCE into CD4^+^ T cells and are thus decisive for T cell activation (12). To quantify the intracellular Ca^2+^ activity ([Ca^2+^]i) and SOCE from control and UA treated CD4^+^ T cells, Fura-2 fluorescence was determined. CD4^+^ T cells were activated for 3 days in the presence of plate-bound anti-CD3 and anti-CD28 (1:2 ratio) and in the presence or absence of UA (5 - 50 µM). The activated cells were incubated with Fura-2 for 30 minutes in standard HEPES and washed once with standard HEPES. [Ca^2+^]i was measured first in standard HEPES, which was subsequently replaced by Ca^2+^-free HEPES. Intracellular Ca^2+^ stores were depleted by addition of sarco-/endoplasmic reticulum Ca^2+^ ATPase (SERCA) inhibitor thapsigargin (1 µM) in the nominal absence of extracellular Ca^2+^. The subsequent re-addition of extracellular Ca^2+^ was followed by a sharp increase of [Ca^2+^]i. No significant difference was observed in slope and peak of the [Ca^2+^]i when CD4^+^ T cells were treated with 5 µM UA, though both parameters tended to be lower compared with untreated control cells. Both, slope and peak of the [Ca^2+^]i were significantly lower in 10 µM UA treated cells than in control cells (Fig. 1B). Furthermore, increasing the UA concentrations (20 µM and 50 µM) significantly decreased SOCE, when compared with untreated control cells (Fig. 1).

**Fig.1:**
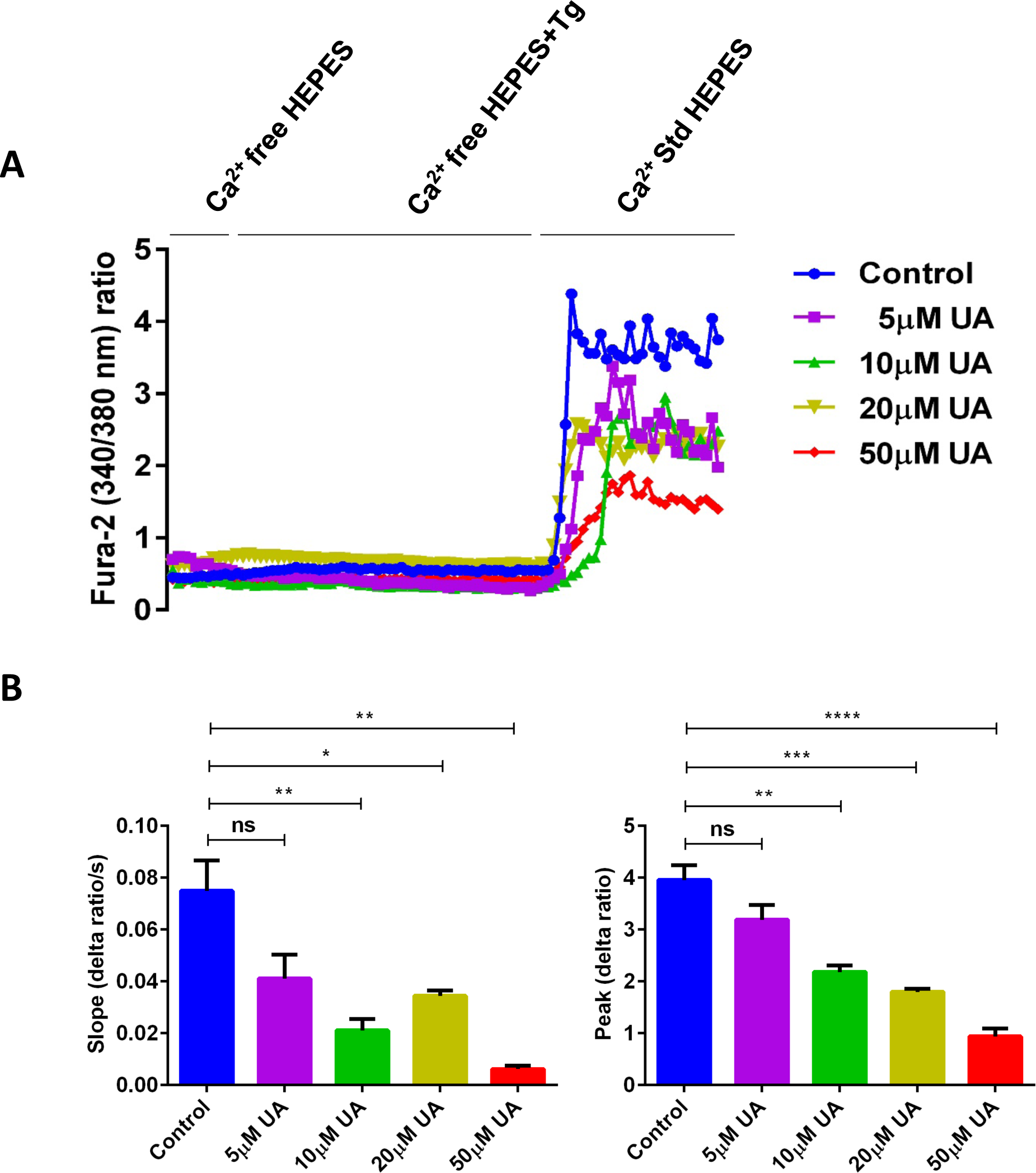
UA treatment significantly decreased SOCE in activated CD4^+^ T cells. A. Representative tracings showing the 340/380 nm fluorescence ratio reflecting cytosolic Ca^2+^ activity in Fura-2, AM loaded activated (plate bound anti-CD3 and anti-CD28) CD4^+^ T cells incubated for 72 hours without and with different concentration of (5 - 50 µM) UA followed by subsequent exposure to Ca^2+^-free HEPES, additional exposure to sarcoendoplasmatic Ca^2+^ ATPase (SERCA) inhibitor thapsigargin (1 µM) and re-addition of extracellular Ca^2+^ (Ca^2+^ Std HEPES). B. Arithmetic means ± SEM (n = 4) of the slope (left) and peak (right) of the fluorescence ratio change following re-addition of extracellular Ca^2+^ in CD4^+^ T cells incubated for 72 hours without (blue bars) and with 5 µM UA (purple bars), 10 µM UA (green bars), 20 µM UA (yellow bars), and 50 µM UA (red bars). *(*p* < 0.05), **(*p* < 0.01), ***(*p* < 0.001), ****(*p* < 0.0001) indicates statistically significant difference.

### UA downregulates the expression of Orai1 and STIM1/2

In preceding experiments, we demonstrated that SOCE (both peak and slope) was significantly downregulated after 10 µM UA. Thus, utilizing qRT-PCR, we further explored whether UA influences Orai1 and/or STIM1/2 transcript levels in CD4^+^ T cells. As illustrated in Fig. 2A, treatment of CD4^+^ T cells with 10 µM UA for 72 hours significantly decreased Orai1 and STIM1/2 mRNA levels. Western blotting was employed to assess, whether the effect of UA on transcript levels was paralleled by similar effects on protein abundance. UA treatment indeed significantly decreased Orai1 protein and STIM1/2 protein expression (Fig. 2B&C).

**Fig.2:**
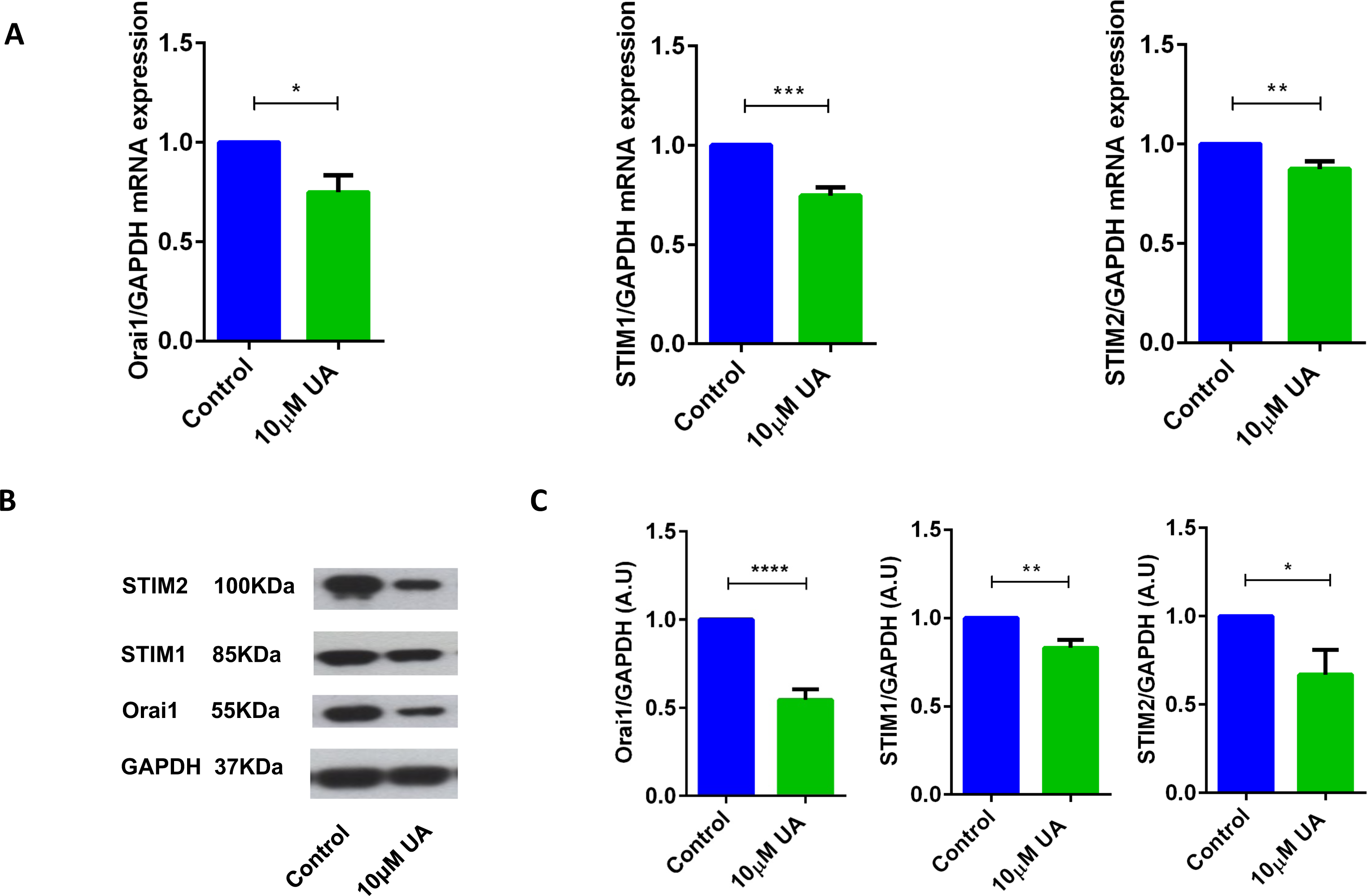
UA significantly decreased Orai1 and STIM1/2 transcripts and protein abundance in CD4^+^ T cells. A. Arithmetic means ± SEM (n = 4 - 7) of Orai1/GAPDH, STIM1/GAPDH and STIM2/GAPDH transcript levels in CD4+ T cells following a 72 hour incubation without (blue bars) and with (green bars) 10 µM UA. *(*p* < 0.05), **(*p* < 0.01), ***(*p* < 0.001) indicates statistically significant difference. B. Original Western blots of STIM1/2 as well as of Orai1 with UA treatment. C. Arithmetic means ± SEM (n = 4 - 7, right panels) of Orai1/GAPDH and (D) STIM2/GAPDH protein abundance in CD4+ T cells following a 72 hour incubation without (control, blue bars) and with (UA, green bars) 10 µM UA. *(*p* < 0.05), **(*p* < 0.01), ****(*p* < 0.0001) indicates statistically significant difference.

### UA treatment augments the miR-10a-5p expression in CD4^+^ T cells

Various metabolites and natural plant products are involved in the regulation of miRNAs biogenesis (32). Previously, we have shown that miR-15b was involved in the regulation of SOCE/STIM2 when induced by green tea polyphenol EGCG (31). Bioinformatics analysis (<u>www.microrna.org</u>, <u>www.targetscan.org</u>, <u>www.mirbase.org)</u> suggested that in addition to miR-15b, several other miRNAs such as miR-10a-5p, miR-29, miR-146 could also modify Ca^2+^ regulating proteins such as Orai/STIM. Thus, we explored whether UA influences miR-10a-5p abundance. To this end, we measured the miR-10a-5p expression in murine CD4^+^ T cells after treatment with UA (5, 10, 20, 50 µM) utilizing the miR-qRT-PCR method. As a result, treatment of CD4^+^ T cells with 10 µM UA was followed by a marked and highly significant increase of miR-10a-5p abundance (Fig. 3). Several other miRNAs such as miR-15b, miR-29a-3p, miR-146a-5p and miR-155-5p were also upregulated after 10 µM UA treatment (Suppl. Fig. 1).

**Fig.3:**
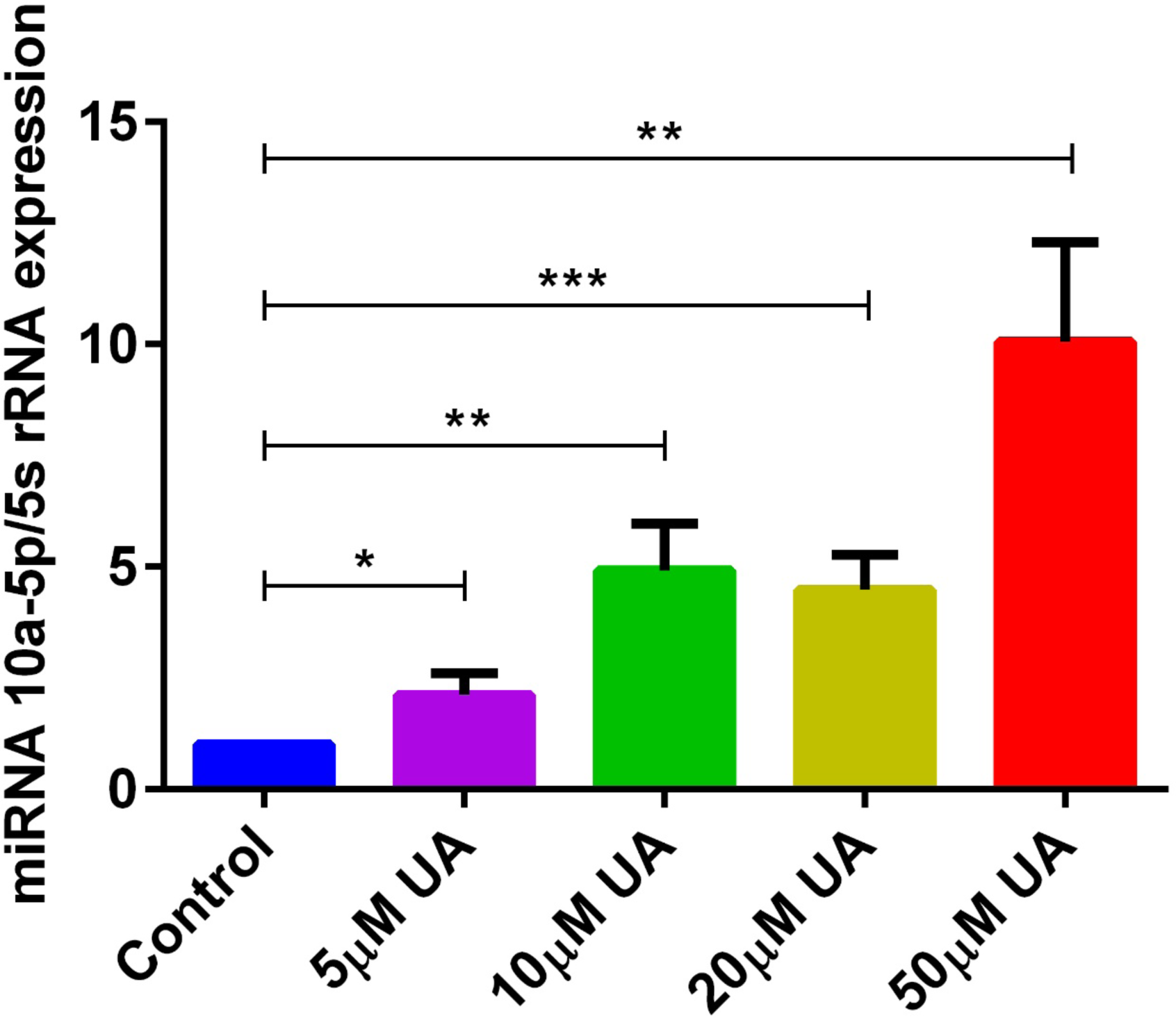
UA treatment significantly increased miR-10a-5p expression in CD4+ T cells. Arithmetic means ± SEM (n = 7) of miR-10a-5p over 5S rRNA transcript levels in CD4+ T cells following a 72 hour incubation without (blue bar) and with 5 µM UA (purple bar), 10 µM UA (green bar), 20 µM UA (yellow green), and 50 µM UA (red bar). *(*p* < 0.05), **(*p* < 0.01), ***(*p* < 0.001), indicates statistically significant difference.

### miR-10a-5p gain and loss inversely affects Orai1 and STIM1/2 transcript and protein levels

Bioinformatics analysis revealed that miR-10a-5p has a strong binding site in the 3’untranslated region (3’UTR) of Orai1, and is thus a potential regulator of Ca^2+^ entry (Fig. 4A). To confirm whether miR-10a-5p influenced Orai1 and/or STIM1/2 transcription, we transfected CD4^+^ T cells with negative mimic and miR-10a-5p mimic and measured the Orai1 and STIM1/2 transcript levels. The qRT-PCR data suggested a marked and highly significant downregulation of both Orai1 and STIM1/2 transcript levels following miR-10a-5p overexpression (Fig. 4B). The decrease of transcript levels was paralleled by similar alterations of protein levels. As apparent from Western blotting, miR-10a-5p overexpression was followed by downregulation of Orai1 and STIM1/2 protein abundance (Fig. 4C&D). Overexpression of miR-10a-5p thus decreased Orai1 and STIM1/2 expression both at transcripts and protein levels. When miR-10a-5p was inhibited in CD4^+^ T cells, a significant increase of both Orai1 and STIM1/2 was observed at transcript levels (Fig. 4E) and protein levels (Fig. 4F&G).

**Fig.4:**
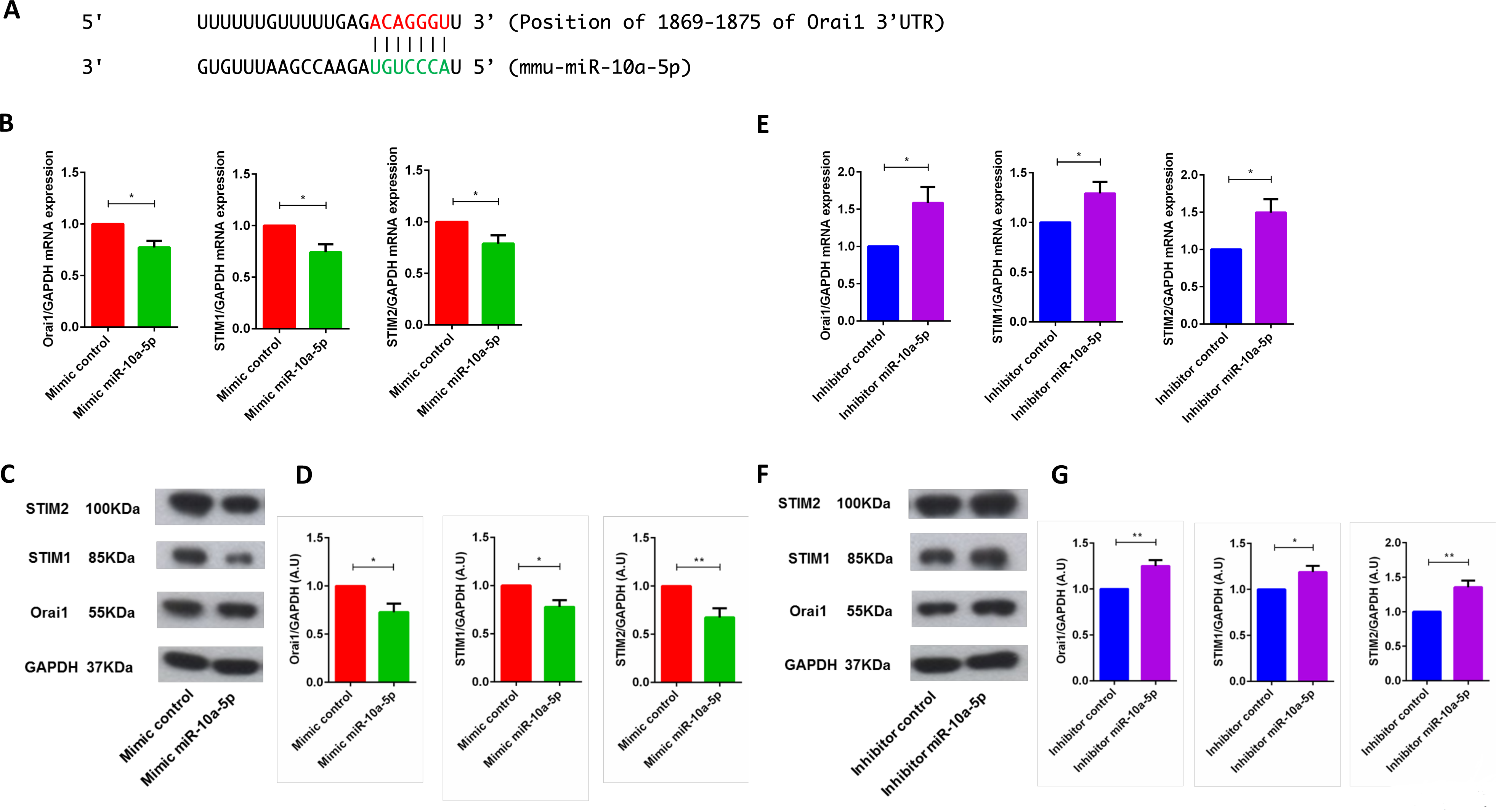
miR-10a-5p mimic over-expression and inhibition significantly decreased and increased Orai1 and STIM1/2 transcript levels and protein abundance in CD4+ T cells, respectively. **A.** Cartoon is showing the miR-10a-5p matches with Orai1 3’-untranslated region (3’-UTR) with seed sequence. **B-D.** Arithmetic means ± SEM (n = 4) of (**B**) Orai1/GAPDH, STIM1/GAPDH and STIM2/GAPDH transcript levels in mimic control (red bars) and miR-10a-5p mimic (green bars) transfected CD4+ T cells. (**C**) Original Western blots (left panels) and (**D**) arithmetic means ± SEM (n = 5, right panels) of Orai1/GAPDH, STIM1/GAPDH and STIM2/GAPDH protein abundance in CD4+ T cells in mimic control (red bars), miR-10a-5p mimic (green bars) transfected CD4+ T cells. *(*p* < 0.05), **(*p* < 0.01) indicates statistically significant difference. **E-G.** Arithmetic means ± SEM (n = 5) of (**E**) Orai1/GAPDH, STIM1/GAPDH and STIM2/GAPDH transcript levels in inhibitor control (blue bars) and miR-10a-5p inhibitor (purple bars) transfected CD4+ T cells. (**F**) Original Western blots (left panels) and (**G**) arithmetic means ± SEM (n = 5, right panels) of Orai1/GAPDH, STIM1/GAPDH, and STIM2/GAPDH protein abundance in CD4+ T cells in inhibitor control (blue bars), miR-10a-5p inhibitor (purple bars) transfected CD4+ T cells. *(*p* < 0.05), **(*p* < 0.01) indicates statistically significant difference.

### miR-10a-5p overexpression and inhibition inversely influence SOCE in CD4^+^ T cells

To examine, whether the downregulation of Orai1 and STIM1/2 expression following *miR-10a-5p* overexpression was paralleled by a similar decrease of SOCE, both control mimic and *miR-10a-5p* mimic transfected CD4^+^ T cells were activated for 3 days in the presence of plate-bound anti-CD3 and anti-CD28 (1:2 ratio). Ca^2+^ entry was measured at day 3 after transfection of miR-10a-5p overexpression using miRNAs mimic. The activated cells were again loaded with Fura-2 for 30 minutes in standard HEPES and washed once with standard HEPES. [Ca^2+^]i was measured first in standard HEPES, which was subsequently replaced by Ca^2+^-free HEPES. In a next step the intracellular Ca^2+^ stores were depleted by addition of thapsigargin (1 µM) in the nominal absence of extracellular Ca^2+^. The subsequent re-addition of extracellular Ca^2+^ was followed by a sharp increase of [Ca^2+^]i. Both, slope and peak of the [Ca^2+^]i increase were significantly lower in miR-10a-5p mimic transfected than in control mimic transfected cells (Fig. 5A&B). Thus, our data suggest indeed that overexpression of miR-10a-5p contributes to the downregulation of Orai1 and STIM1/2 expression following UA treatment. Conversely, inhibition of miR-10a-5p augments significantly both slope and peak of the [Ca^2+^]i increase following Ca^2+^ re-addition (Fig. 5C&D). Our gain-of-function and loss-of-function data suggested that indeed miR-10a-5p is a powerful regulator of SOCE.

**Fig.5:**
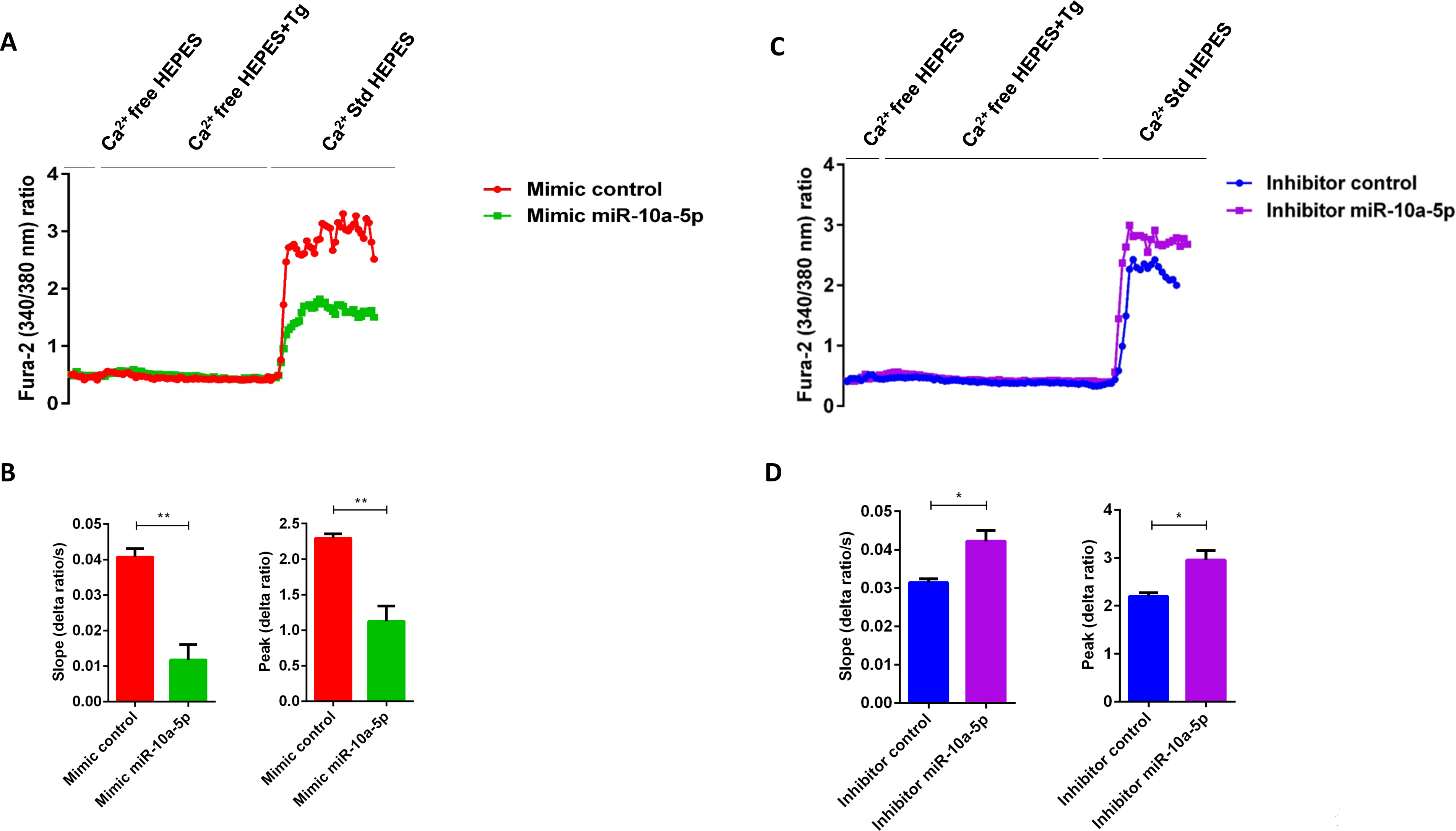
miR-10a-5p overexpression and inhibition lead to loss-of-function and gain-of-function of SOCE in activated CD4^+^ T cells, respectively. A. Representative tracings showing the 340/380 nm fluorescence ratio reflecting cytosolic Ca^2+^ activity in Fura-2/AM loaded negative mimic control (red), and miR-10a-5p mimic (green) transfected CD4+ T cells following exposure to Ca^2+^-free HEPES, additional exposure to thapsigargin (1 µM) and re-addition of extracellular Ca^2+^ (Ca^2+^ Std HEPES). B. Arithmetic means ± SEM (n = 3) of the slope (left) and peak (right) of the fluorescence ratio change following re-addition of extracellular Ca^2+^ in negative mimic control (red bars) and miR-10a-5p mimic (green bars) transfected CD4+ T cells. **(*p* < 0.01) indicates statistically significant difference. C. Representative tracings showing the 340/380 nm fluorescence ratio reflecting cytosolic Ca^2+^ activity in Fura-2/AM loaded inhibitor control (blue), and miR-10a-5p inhibitor (purple) transfected CD4+ T cells following exposure to Ca^2+^-free HEPES, additional exposure to thapsigargin (1 µM) and re-addition of extracellular Ca^2+^ (Ca^2+^ Std HEPES). D. Arithmetic means ± SEM (n = 3) of the slope (left) and peak (right) of the fluorescence ratio change following re-addition of extracellular Ca^2+^ in inhibitor control (blue bars), and miR-10a-5p inhibitor (purple bars) transfected CD4+ T cells. *(*p* < 0.05) indicates statistically significant difference.

### Effect of UA on cell proliferation

As stimulation of SOCE is involved in the signalling triggering T-cell proliferation (25), cell proliferation was quantified using the dye CFSE. As illustrated in Fig. 6, cell proliferation was significantly decreased in the presence of 10 µM UA.

**Fig.6:**
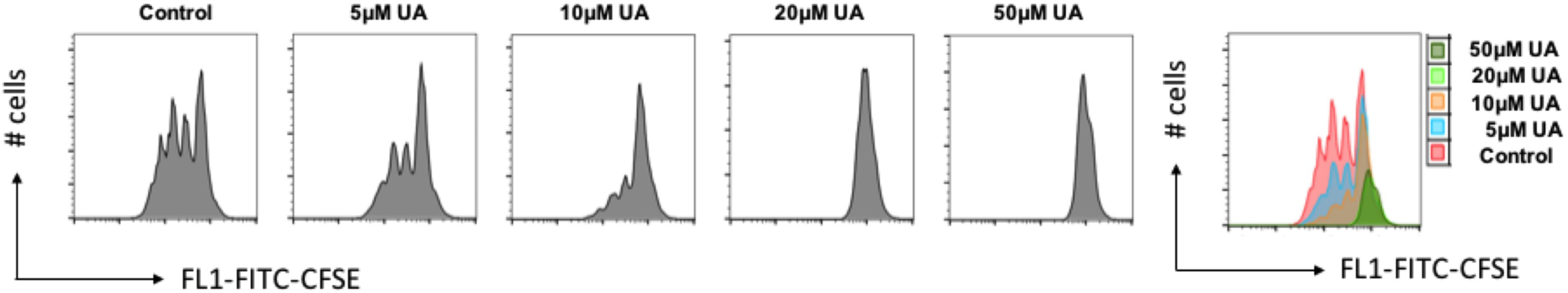
UA decreased cell proliferation of CD4^+^ T cells. A. WT CD4+ T cells were stained with CFSE dye before activation with anti-CD3/anti-CD28 and cultured in the presence of (5 - 50 µM) UA for 3 days. Cell proliferation was measured by flow cytometry. Data are representative for 3 independent experiments. X-axis represents the CFSE dye whereas y-axis represents cell numbers (# no. of cells). B. Overlays plot of cell proliferation with different concentrations of UA. X-axis represents the CFSE dye whereas y-axis represents cell numbers (# no. of cells).

## DISCUSSION

The polyphenolic compounds are known to control inflammation, angiogenesis, drug and radiation resistance (1). These findings have been further corroborated by the observations that Ellagitannin-rich food have beneficial effects on inflammatory bowel disease and other inflammatory diseases (2). However, the bioavailability of these compounds (ellagitannins and ellagic acid) is limited and the compounds must be metabolized by the gut microbiota to produce bioactive molecules that can be easily absorbed (7). Ellagic acids (by the help of gut bacteria) are converted into different Urolithins (A-D) (7). Gut metabolites such as Urolithins can directly activate adaptive immune T cells (8–11). In this report, we reveal that UA can suppress store operated Ca^2+^ entry by modulating Orai/STIM regulating miRNAs, thus affecting Ca^2+^ sensitive cellular functions including CD4^+^ T cell proliferation.

High levels of intracellular Ca^2+^ are necessary to maintain numerous functions of T cells such as the interaction between a T-cell and antigen-presenting cell (APC) that leads to formation of the specialized contact surface known as the immunological synapse and activation of different transcription factors (12, 18, 22, 33, 34). Several hours of oscillating Ca^2+^ influx are required to complete the T-cell activation program, which involves expression of a large number of activation-associated genes (25). Our observations describe for the first time that bacterial metabolite product UA is a negative regulator of store operated Ca^2+^ entry into murine CD4^+^ T cells, an effect paralleled by downregulation of Orai1/STIM1/2 expression. Thus, the present observations also uncover a completely novel mechanism accounting for the effect of UA on SOCE, i.e. the upregulation of miR-10a-5p, which in turn downregulates Orai1 and STIM1/2 transcript and protein levels as well as SOCE. Thus, UA changes the post-transcriptional machinery of the key players undertaking SOCE in CD4^+^ T cells, i.e. Orai1 and STIM1/2.

Emerging evidence indicates that UA is involved in the regulation of inflammatory pathways, cell cycle and cell death (1, 35, 36). Some studies further suggested that UA is also involved in controlling the growth of cancer cells, and could thus be used as anti-cancer agent (1, 35). It is tempting to speculate that UA interferes similarly with tumor cell proliferation by downregulating SOCE.

Recently, we have demonstrated that miRNAs processing protein *dicer* is involved in the regulation of SOCE in CD4^+^ T cells (30). Thus, identifying the role of individual miRNAs which could be involved in the regulation of Ca^2+^ pathways open a new avenue to therapeutic intervention. Using bioinformatics tools, we found that miR-10-5p could regulate Orai1 proteins thus modifying STIM1/2 expression. Previous studies have reported that miR-10a-5p is involved in the development and function of regulatory T cells (Tregs) (37). Our results reveal that miR-10a-5p was upregulated after treatment with UA in T cells. Thus, miR-10a-5p appears to be involved in the regulation of Ca^2+^ entry and thus Ca^2+^ sensitive cellular functions such as gene expression, proliferation, cell motility and cytokine expression. However, a role of other miRNAs or further signalling pathways cannot be excluded.

In conclusion, the present observations reveal a completely novel role of gut bacterial metabolite UA in the regulation of Ca^2+^ entry into CD4^+^ T cells leading to suppression of activation of CD4^+^ T cell activation. UA upregulates the expression of miR-10a-5p which in turn decreases SOCE by downregulating Orai1 and STIM1/2 expression. Thus, our results suggest that upregulation of miR-10a-5p by UA restrains store operated Ca^2+^ entry in murine CD4^+^ T cells and UA could be used a natural immune-suppressant during various inflammatory disorders including inflammatory bowel disease.

## MATERIALS AND METHODS

### Mice

Naïve CD4^+^ T Cells were isolated from C57BL/6 mice (male and female) between 8 - 16 weeks of age. All the animals were kept in standard housing conditions with 12 ± dark/light cycle and fed on chow diet. All experiments were performed according to the EU Animals Scientific Procedures Act and the German law for the welfare of animals. All procedures were approved by the authorities of the state of Baden-Württemberg, Germany.

### Naïve CD4^+^ T cell isolation and culture

Naïve CD4^+^ T cells were purified from spleen and lymph nodes of mice C57BL/6 using the MagniSort^®^ Mouse naïve T cell Enrichment kit (# 8804-6824-74, eBioscience, USA) as described by the manufacturer. Purified naïve CD4^+^ T cells were cultured in plate-bound anti-CD3 (#16-0031-85, eBioscience, USA)/anti-CD28 (#16-0281-85, eBioscience, USA) Abs at a 1:2 ratio (1µg/ml anti-CD3 and 2µg/ml anti-CD28) in the presence or absence of 5 - 50 µM UA (#1143-70-0, Santa Cruz Biotechnology, USA) for 3 days.

### Intracellular Calcium measurement

Intracellular Ca^2+^ activity was measured using Fura-2-AM (#F1221, Molecular Probes, USA). Fluorescence measurements were performed using an inverted light incidence fluorescence phase-contrast microscope (Axiovert 100, Zeiss, Germany). Cells were excited alternatively at λ = 340 or 380 nm and the light deflected by a dichroic mirror into either the objective (Fluar 40×/1.30 oil, Zeiss, Germany) or a camera (Proxitronic, Germany). Emitted fluorescence intensity recorded at λ = 505 nm and data were acquired by using specialized computer software (Metafluor, Universal Imaging, USA) (38).

Activated T cells (3 days) were loaded with 2 µM Fura-2, AM for 30 min at 37°C in a CO2 incubator. To measure SOCE, changes in cytosolic Ca^2+^ activity ([Ca^2+^]i) were monitored following depletion of the intracellular Ca^2+^ stores. In short, [Ca^2+^]i was measured using Ca^2+^ containing standard HEPES buffer [125mM/L NaCl, 5mM/L KCl, 1.2 mM/L MgSO4*7H2O, 32.2 mM/L HEPES, 2mM/L Na2HPO4*2H2O, 5mM/L Glucose, 1mM/L CaCl2*2H2O; pH=7.4] for 2 minutes and then changed to Ca^2+^-free HEPES buffer [125mM/L NaCl, 5mM/L KCl, 1.2 mM/L MgSO4*7H2O, 32.2 mM/L HEPES, 2mM/L Na2HPO4*2H2O, 5mM/L Glucose, 0.5 mM/L EGTA; pH=7.4] for 3 minutes. In the absence of Ca^2+^, the intracellular Ca^2+^ stores were depleted by inhibition of the sarcoendoplasmatic Ca^2+^ ATPase (SERCA) by 1 µM thapsigargin (#67526-95-8, Sigma, Germany) and [Ca^2+^]i was measured for another 5 minutes. In the following, Ca^2+^ containing HEPES buffer was added for 5 minutes, which allowed assessing the SOCE.

### Transfection of CD4^+^ T cells by miR-10a-5p

Naïve CD4^+^ T cells were seeded on a coated 24-well plate. Naïve T cells were transfected with miR-negative mimic (#479903), miR-10a-5p mimic (#471928), miR-10a-5p inhibitor (#4100036) (Exiqon, Denmark) using DharmaFECT3 (#T-2003-01, Dharmacon, USA) as recommended by manufacture’s guidelines. Briefly, naïve T cells were prepared in antibiotic free cell buffer and 0.75×10^6^ - 1×10^6^ cells per well cultured in the presence of 500 µl of R-10 medium (RPMI 1640 (#61870-010, Life technologies, USA) medium supplemented with 10% fetal bovine serum (#10270-106, life technologies, USA), 1% L-Glutamine (#G7513 200mM solution, Sigma, Germany), 1% penicillin/streptomycin (#P4333, Sigma, Germany) and 0.1% 2-Mercaptoethanol (#31350-010, Life technologies, USA). Whilst plating the cells, 2 µl of 50 µM stock concentration of non-targeting miRNA-negative mimic, miR-10a-5p-mimic or miR-10a-5p-inhibitor were added to 8 µl of antibiotic free RPMI1640 medium and the cells incubated for 5 minutes in tube 1, respectively. In tube 2, 0.5 µl of DharmaFECT3 was added to 9.5 µl of antibiotic free RPMI1640 medium. The content of tube 1 was added to tube 2 and incubated for additional 20 minutes. After 20 minutes of incubation, the reaction mixture from tube 2 was added to corresponding wells to control and miR-10a-5p-mimic or miR-10a-5p-inhibitor wells. Cells were further incubated for additional 72 hours and used for qRT-PCR, immunoblotting, and determination of SOCE.

### mRNA and miRNA qRT-PCR

Total RNA including miRNAs was extracted from CD4^+^ T cells using miRNAeasy Kit (#217004, Qiagen, Germany). The mRNA (1 µg) and miRNAs (100 ng) were separately reverse transcribed using Superscript III First-Strand synthesis system (#18080-51, Invitrogen, Germany) and miRNA universal cDNA synthesis kit II (#203301, Exiqon, Denmark) for reverse transcript PCR (RT-PCR) and subsequent real-time quantitative PCR (qRT-PCR). Detection of gene expression was performed with KapaFast-SYBR Green (#KAPBKK4606, Peqlab, Germany) and measurements were performed on a BioRad iCycler iQ^TM^ Real-Time PCR Detection System (Bio-Rad Laboratories). The relative expression levels of mRNAs were normalized to that of *GAPDH*, whereas the relative expression levels of miRNAs were normalized to that of 5S rRNA. The following murine primers were used to detect *Orai1*, *STIM1*, and S*TIM2* expression (39).

Orai1-F 5’-CCTGGCGCAAGCTCTACTTA-3’

Orai1-R 5’-CATCGCTACCATGGCGAAGC-3’

STIM1-F 5’-ATTGTGTCGCCCTTGTCCAT-3’

STIM1-R 5’-TGGGTCAAATCCCTCTGAGAT-3’

STIM2-F 5’-TGTCTGTGTCAAGTTGCCCT-3’

STIM2-R 5’-TGTCTGGCACTTCCCATTGT-3’

GAPDH-F 5’-CGTCCCGTAGACAAAATGGT -3’

GAPDH-R 5’-TTGATGGCAACAATCTCCAC-3’

For amplification of different miRNAs, hsa-miR-10a-5p LNA™ PCR primer set (#204778, Exiqon, Denmark), hsa-miR-15b-5p LNA™ PCR primer set (#204243), hsa-miR-29a-3p LNA™ PCR primer set (#204698), hsa-miR-146a-5p LNA™ PCR primer set (#204688), mmu-miR-155-5p LNA™ PCR primer set (#205930), and reference 5S rRNA primer set (#203906) were used and the reaction was set up as recommended by Exiqon or described earlier (40–42).

### Immunoblotting

CD4^+^ T cells were activated in presence of anti-CD3 (1µg/ml)/anti-CD28 (2µg/ml) and treated with 10 µM UA. After 72 hours of activation and treatment, CD4^+^ T cells were washed once with PBS, counted and equal amounts cells were taken for cell lysis using H2O and 2×Lammelli’s Buffer. Proteins were denatured at 95^0^C for 5-10 minutes and stored at -20^0^C. Sample proteins were loaded on 8% or 10% gel depending on protein size and run at 80 V until crossing of stacking gel then voltage was increased to 120 V during the separation phase and total gel run for 90-100 minutes. Proteins were electro-transferred onto PVDF membranes. Membranes were probed with the indicated primary antibodies for Orai1 (1:1000; #13130-1-AP, Proteintech, United Kingdom), STIM1 (1:1000; #5668S, Cell Signaling Technology), STIM2 (1:1000; #4917S, Cell Signaling Technology) and GAPDH (1:2000; #5174S, Cell Signaling Technology), followed by HRP-conjugated secondary antibodies (1:1000; #7074P2, Cell Signalling Technology, Germany). Membranes were washed and visualized with enhanced chemiluminescent HRP substrate (#R-03031-D25 and R-03025-D25, advansta, USA). Data were analysed by ImageJ software (<u>https://imagej.nih.gov/ij/)</u>.

### CFSE staining

The proliferation of CD4^+^ T cells was detected by CellTrace^TM^ CFSE Cell Proliferation Kit (#C34554, eBioscience, USA). Briefly, cells were washed with PBS (#D8537, Sigma, Germany) once, stained with CellTrace^TM^ CFSE (1:1000 dilution) and re-suspended gently, incubated at 37^0^C for 15 minutes in the dark, then washed with R-10 medium twice and activated as described in *CD4^+^ T cell isolation and culture* in Materials and Methods above. After 72 hours, cells were collected to perform the flow cytometry. Data were analysed by Flowjo software (FLOWJO LLC, USA).

### Statistics

Data are provided as means ± SEM, n represents the number of independent experiments. All data were tested for significance using unpaired Student’s t-test or ANOVA. Data were analysed by Excel 2010 or GraphPad Prism Software, USA. P value ≤0.05 was considered statistically significant.

## AUTHOR CONTRIBUTIONS

SZ, TAM, HC, LP, MSS and YS performed the research and analysed the data. AC, FL and YS designed the study, supervised the project and wrote the manuscript. All authors edited and approved the final manuscript.

## ACKNOWLEDGEMENTS

The Research is supported by Deutsche Forschungsgemeinschaft research grant (F.L.). MSS is supported by ZUK63 Tübingen university fellowship and Fortüne grant (2426-0-0). TAM is supported by DAAD PhD fellowship, HC is supported by China Scholarship Council and LP is supported by Brigitte-Schlieben-Lange-Programme. Both SZ and AC are supported by China Agricultural Research System (CARS-42-17). We also acknowledge the support by “The Open Access Publishing Fund Tübingen University”.

## DISCLOSURE STATMENT

The authors of this manuscript state that they do not have any financial conflict of interests.

## FIGURE LEGENDS

**Supp. FIG 1.**
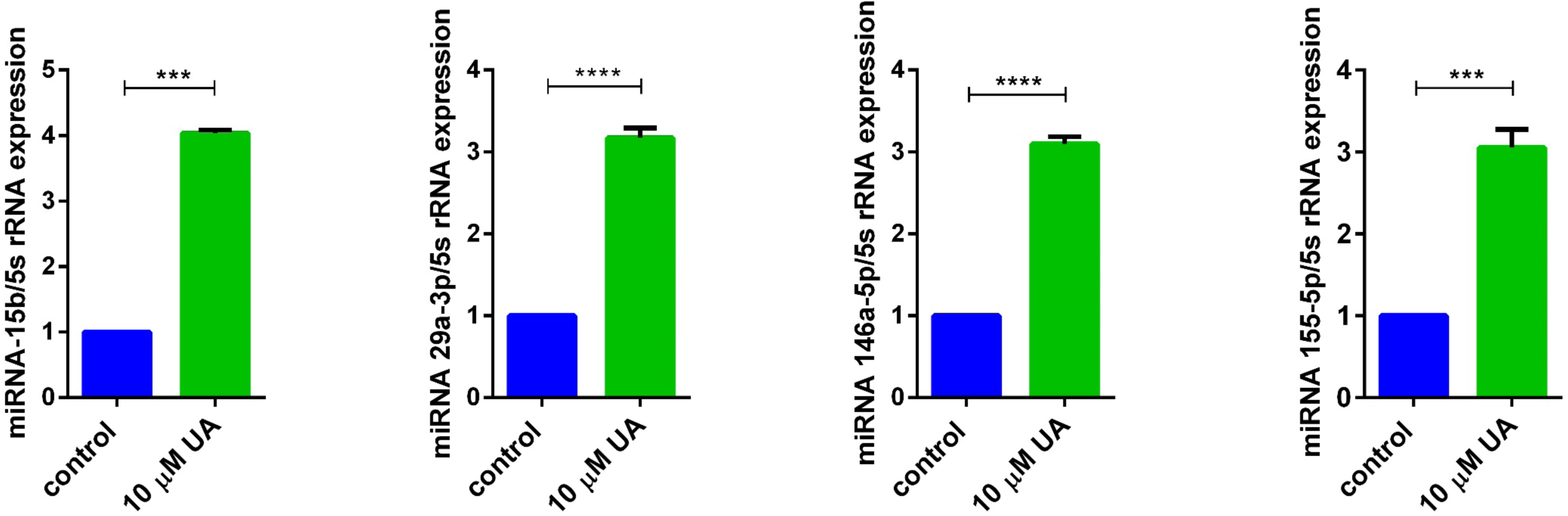

